# Dual scRNA-Seq analysis reveals rare and uncommon parasitized cell populations in chronic *L. donovani* infection

**DOI:** 10.1101/2022.07.26.501600

**Authors:** Konstantinos Karagiannis, Sreenivas Gannavaram, Chaitenya Verma, Parna Bhattacharya, Hira L Nakhasi, Abhay Satoskar

## Abstract

Although phagocytic cells are documented targets of *Leishmania* parasites, it is unclear whether these parasites can infect other cell types. In this study, we describe a computational approach that exploits scRNA-seq to simultaneously analyze the transcriptomic signatures of the host cell and to identify rare and uncommon cells that harbor *Leishmania donovani* in the spleen and bone marrow. Individual cells were annotated as parasitized based on the presence of *L. donovani* transcripts that were detected with high accuracy. This unbiased approach allowed identification of heterogenous parasitized cell populations that cannot be detected by conventional methods. Consistent with previous studies, analysis of spleen cells isolated from *L. donovani* infected mice revealed inflammatory monocytes as the dominant parasitized cells. In addition, megakaryocytes, basophils, and NK cells were found to be the rare cells infected in the spleen. Unexpectedly, hematopoietic stem cells (HSCs), not known to be phagocytic, were the dominant cells parasitized cell in the bone marrow. In addition, eosinophils, megakaryocytes, and basal cells were the rare bone marrow cells found to be infected. scRNA-seq analysis revealed known phagocytic receptors Fc_γ_R and CD93 are expressed on HSCs. *In vitro* studies using purified HSCs showed that these cells can phagocytize *L. donovani*. Parasitized HSCs were also detectable in the bone marrow of mouse infected with *L donovani*.. This unbiased dual scRNA-seq approach enables identification of rare and uncommon parasitized cells that could be involved in pathogenesis, persistence, and protective immunity. Further, such approach could be used to study pathogenesis of other infectious agents.

## Introduction

Protozoan organisms belonging to *Leishmania* species are the parasites of the mammalian reticuloendothelial system (WHO, 2021). Depending on the species, *Leishmania* parasites cause three distinct clinical manifestations, i.e., cutaneous (CL), mucocutaneous (MCL) and visceral leishmaniases (VL). WHO estimates indicate that about 1,200,000 cases of CL and 700,000 cases of CL occur annually (WHO, 2021). VL is the most clinically severe form that often results in death if untreated and is accompanied by hepato-splenomegaly, anemia, and fibrosis and dyserythropoiesis in bone marrow (Goto et al., 2017, Saleem et al., 1991). Only a small fraction of the infections are documented to result in symptomatic VL in humans while the majority remain as asymptomatic carriers (WHO, 2021). The immunological mechanisms dominated by the Th-1 polarized response characterized by pro-inflammatory cytokines IFN-γ, IL-12 and TNF have been shown to clear the parasites in asymptomatic VL (Kaye et al., 2004, Engwerda et al., 2004), yet asymptomatic *L. donovani* infections and PKDL are intensely studied since they are considered to facilitate disease transmission (Hirve et al., 2016). Therefore, it begs the question what are the host cell reservoirs of the parasite in chronic infections? The complex interactions of host and parasitic factors that determine the pathophysiology, clinical variability, susceptibility/resistance and chronicity are intensely studied towards achieving the goals of control and elimination of VL (Alvar et al., 2021, Poulaki et al., 2021).

The advent of high throughput methods such as scRNA-Seq matched by new computational analyses enabled probing of such diverse problems as hematopoiesis (Zheng et al., 2018), embryonic development (Chu et al., 2016, Haniffa et al., 2021), and construction of single cell atlases for various cancers (Wagner et al., 2019, Neftel et al., 2019). Similar methods have been applied to study parasitic diseases in terms of the tripartite interactions between host cells, parasites and vectors. Application of scRNA-Seq analysis revealed the hidden diversity of transcriptional signatures in different life stages of *Plasmodium* parasites associated with pathogenicity (Read et al., 2018), sexual commitment (Poran et al., 2017), asynchronous development and maturation of sporozoites (Bogale 2021). Single cell epigenomics and scRNA-Seq tracked the developmental dynamics of Th1 and Tfh cells that influence the emergence of CD4+ T memory cells in malaria (Lonnberg et al., 2017, Soon et al., 2020). A convergent transcriptional profile in atypical B cells has been identified in malaria, HIV and autoimmune disease (Holla et al., 2021). A comprehensive functional genomics analysis by scRNA-Seq on various species of *Plasmodium* parasites and parasite-infected cells produced an interactive Malaria Cell Atlas that has been hailed as a roadmap for malaria research (Notzel et al., 2021, Winzeler 2019, Howick et al., 2019).

Similar to studies in *Plasmodium* parasites, scRNA-Seq has been applied to track genetic hybrids of *Leishmania* following exposure to DNA damage stress that enhanced efficiency of in vitro hybridization of the *L. tropica L. donovani, L. infantum*, and *L. braziliensis*, a capacity to generate intra- and interspecific hybrids (Louradour et al., 2022). Numerous studies with various species of *Leishmania* applied bulk RNA-seq methods to analyze drug resistance (Perea-Martínez et al., 2022), RNA interactome (Kalesh et al., 2022), host-parasite interactions (Dillon et al., 2015), and characterization of the developmental stages of *L. major* in the sand fly (Inbar et al., 2017). Dual RNA-Seq analysis that targeted both host and parasite transcriptomes by bulk RNA-seq in liver and spleen of *L. donovani* and *L. infantum* infected mouse models showed potential differences in the pathophysiology in the two infections (Forrester et al., 2022). Transcriptional analysis using in silico cell type deconvolution identified a gene expression signature that was conserved in VL cohorts in India, Ethiopia and Brazil (Dirkx et al., 2022). scRNA-Seq of the cells isolated from *L. major* infected mice identified recruited and skin resident heterogenous populations (Venugopal et al., 2022).

Development of a *Leishmania* Atlas analogous to Malaria Cell Atlas would facilitate unprecedented investigations into such intractable questions as the parasite persistence, drug resistance, determinants of protective immunity, and pathogenesis. Despite rapid advances in computational tools, single cell based RNA-seq methods have never been applied to analyze both host and parasite transcripts simultaneously. Analysis of host and parasite transcripts at single cell level would enable identification of novel cell types by virtue of cell annotation by transcriptomic analysis. The conventional methods as flow cytometry are limited in their application since they can only detect cells for which markers are available. Towards this goal, we describe a novel dual sc-RNAseq method that identifies parasitized cells from the spleen and bone marrow of mice chronically infected with *L. donovani* and to analyze transcriptomic signatures from the parasitized and bystander cells to gain deep insights into the pathogenesis of VL. Such an agnostic method can be widely applied to other parasitic infections for which reference transcriptomes are available.

## Methods

### Mouse infection

Wildtype female Balb/C mice of 9-10 weeks age were purchased from Envigo (Indianapolis, IN). Mice were housed and maintained at The Ohio State University (OSU) Laboratory Animal Resources Facilities. Mice were infected intravenously with 1×10^7^ Ds-Red-expressing *L. donovani* amastigotes as described before (Terrazas et al., 2015). Mice (n=2) were euthanized using a CO_2_ chamber at 60 days post infection (dpi) corresponding to chronic infection.

### Sample preparation for single cell suspension

Spleen and bone marrow were aseptically harvested from the mice and a single cell suspension was prepared from spleen by gentle mechanical disruption and passed through a 70μm cell strainer (Cat. 352350; Falcon™, Corning, NY, USA). For the bone marrow cells, femur and tibias were isolated and cut at both ends with scissors, and cells were flushed out from the cavity into RPMI medium-1640 (Cat. 11-875-093; Gibco, Thermo Fisher, Grand Island, NY, USA) supplemented with 10% fetal bovine serum (Cat. 97068-085; Avantor Seradigm, Ballycoolin, Dublin, Ireland), 1% penicillin and streptomycin (Cat. 15140-122; Gibco, Thermo Fisher), and 0.1% β-mercaptoethanol (Cat. 21985-023; Gibco, Thermo Fisher). Cells were passed through a MACS-LS column (Cat no. 130-042-401) to remove the non-targeted cells by using monocyte isolation kit (Cat. 130-100-629, Milteny Biotech; Bergisch Gladbach, Germany). Monocyte enrichment was achieved by depletion of magnetically labeled non-targeted T cells, B cells, NK cells, dendritic cells, erythroid cells, and granulocytes.

### Preparation of libraries and single cell RNA sequencing

Single-cell RNA sequencing was performed using the Chromium (10x Genomics) instruments. Cells in single cell suspension were counted using a hemocytometer, viability was measured by trypan blue (viability: Bone marrow; 88% and Spleen; 97%), and the volume was adjusted to 1000 cells/μl. Cells were processed using the 10x Genomics Chromium Controller and the Chromium Next GEM Single cells 3’ Library Kit v3.1 (PN-1000268; 10x Genomics, Pleasanton, CA, USA) following the standard manufacturer’s protocol. Briefly, cells were loaded into the chromium controller aiming for 10,000 captured cells for library preparation and sequencing. The integrity of the cDNAs was evaluated using Agilent Bioanalyzer High Sensitivity Chip (Agilent Technology) and the quantity was assessed by Qubit assay. Cellular suspension of cDNA library was loaded on Chromium Next GEM Chip G to generate GEM (Gel Bead-in-emulsion) barcoding along with the single cell reagent to create micro-reaction and GEM clean-up by using Chromium Next GEM Chip G Single Cell Kit (PN-1000120; 10x Genomics). In the final step, the Single cell 3’ Gene expression library was amplified by Dual Index Kit TT set A (PN-1000215; 10x Genomics). To prepare the final libraries, amplified cDNA was enzymatically fragmented, and size selected using SPRIselect magnetic beads (Beckman Coulter) followed by ligation of Illumina sequencing adapters. The quality of the final library was assessed using an Agilent Bioanalyzer High Sensitivity chip. Samples were then sequenced on the Illumina NovaSeq 6000 sequencer with a target of (>15,000number of reads/cells i.e.,_paired end reads), yielding a median per-library depth of 27,637 reads per cell.

### Single cell RNA-seq reference

For the single cell analysis, genomic sequences for *Mus musculus* (assembly GRCm38 with accession number GCF_000001635.27) *Leishmania donovani* (L. Don) (assembly ASM22713v2 with accession number GCF_000227135.1) were downloaded from RefSeq and merged into one file. Along with the sequences the genomic annotations “.gtf” files were downloaded. Before merging the genome files into one L. Don annotation file was preprocessed by prepending the gene names with the prefix a “ ldon”. The headers of the gff files were merged manually and the annotation information for L. Don was appended to the end of the mouse annotation file. The merged genome and annotation file were then processed by cellranger mkref (version 5.0.1), using the default parameters, to create the appropriate index of an artificial single genome.

### Single cell RNA-seq read processing

The base calling was performed using cellrange mkfastq (version 5.0.1) using the default parameters. The reads of each of the four samples were processed separately using the cellranger count (version 5.0.1) against the host/parasite merged genome and the corresponding annotation file as described above. The output hits were processed using an internally developed script that separated the hits to the host transcripts from those to the parasite transcripts based on the prefixed names of the latter. The alignments produced by cellranger count were processed with samtools (version 1.3.1) to produce host and parasite specific metrics including the estimated number of cells, the median genes per cell, the total number of genes detected and median UMI counts per cell.

### Single cell RNA-seq population annotation

Count matrices produced by cellranger count were processed using the R package Seurat (v4.0) [Butler et al., 2018, Stuart et al., 2019, Hao et al., 2021]. The hits of *L. donovani* transcripts here detected based on the “ldon_” prefix prepended to the names of the transcripts and were separated from the rest of the hits. *L. donovani* transcript hits were summed, and the collapsed feature named “LDon” was used to annotate the cells. Cells with 1 or more *L*. Don *donovani* feature counts was considered parasitized and cells with 0 LDon feature count were considered non-parasitized. The GRCm38 transcripts counts were used for downstream analysis and were normalized using SCTransform and the mitochondrial mapping percentage as a confounding factor. The two replicates from each tissue were integrated using Seurat’s SCTransform specific integration workflow. The cells were clustered by applying the K-nearest neighbors (KNN) graph based on PCA reduced space, followed by Louvain’s algorithm available in Seruat (v4.0). The cells were projected to a two-dimensional space using the Uniform Manifold Approximation and Projection (UMAP) dimensionality reduction technique. The clusters were annotated using the SCSA [Cao et al., 2020] method (https://github.com/bioinfo-ibms-pumc/SCSA accessed in Aug. 2021) and the markers available at the CellMarker [Zhang et al., 2019] curated database. Infected cells of a type with less than 5 parasitized cells in total were excluded from downstream analysis.

### Tertiary analysis of cell populations

Gene markers of each population were detected by identifying significant differentially expressed genes between one population and the rest of the cells using Wilcoxon test, available in Seurat package, including only genes expressed in at least 10% of the cells of either groups. For the tertiary analysis of parasitized cells, subsets of cells were selected from the populations with majority of the infected cells (HSCs for bone marrow and monocytes for spleen). The subsets were further split between two groups, the “infected” and “non infected” cells. Gene expression analysis was performed and significant differentially expressed genes were identified using Wilcoxon test available in Seurat package including only genes expressed in at least 10% of the cells of either groups. Adjusted p values are reported after bonferoni correction. Gene set enrichment analysis was performed on KEGG database (Kanehsia, 2019) using clusterProfiler package (version 3.18.1) in R (Yu et al., 2012). p-values were adjusted using Benjamini & Hochberg method.

### HSC purification from bone marrow

Mouse HSCs were isolated by negative selection from 5-8 week old female Balb/C mouse bone marrow using EasySep™ mouse hematopoietic progenitor cell isolation kit (Stemcell Technologies) following manufacturer’s instructions. Briefly, 10^8^ bone marrow cells were mixed with isolation antibody cocktail followed by RapidSpheres™. After inserting the tube into the magnet, unbound hematopoietic progenitor cells were collected and washed in complete RPMI medium. The purity of the isolation was verified by flow cytometry.

### Flow cytometric analysis

HSCs isolated from mouse bone marrow were rested for 24h by incubation in 37°C incubator and 5%CO_2_ and infected *in vitro* with transgenic *Leishmania donovani-RFP* parasites for 48h at a ratio of 1:5. Cells were stained and analyzed by flowcytometry using Cytek Aurora and analyzed with FlowJo v10 (Treestar). Anti-Ter-119, anti-CD11b, anti-CD5, anti-CD3, anti-CD45R, and anti-Gr-1, all FITC-conjugated, were added at a concentration of 1:100 for 30 minutes at 4°C. Cells were washed with PBS containing 0.5% BSA. Anti-c-Kit-eFluor450, anti-Sca-1-PE-Cy5, anti-CD150-BV650, anti-CD48-BV510, biotinylated anti-Flk2 detected with streptavidin-BV785, anti-CD16/32-PEDazzle594, anti-CD93-PECy7 all were added at a concentration of 1:100 and incubated at 4°C for 30 minutes.

Bone marrow from Balb/C mice chronically infected with *L. donovani^DsRed^* parasites was isolated, subjected to ACK lysis. Cells were passed through a MACS-LS column to enrich monocytes by depletion of magnetically labeled non-targeted T cells, B cells, NK cells, dendritic cells, erythroid cells, and granulocytes as described above. Cells were stained with anti-Ter-119, anti-CD11b, anti-CD5, anti-CD3, anti-CD45R, and anti-Gr-1, (all FITC-conjugated) antibodies at a concentration of 1:100 for 30 minutes at 4°C. Cells were washed with PBS containing 0.5% BSA. Anti-c-Kit-eFluor450, anti-Sca-1-PE-Cy5 antibodies were added at a concentration of 1:100 and incubated at 4°C for 30 minutes and analyzed by flowcytometry using BD Fortessa and analyzed with FlowJo v10 (Treestar) to identify *L. donovani^DsRed^* infected HSCs in vivo.

## Results

### Analysis of the scRNA-seq data

To detect cells parasitized by *L. donovani* and investigate the impact on their transcriptional profile, single cell RNA-seq analysis was performed on tissue samples collected from a chronically infected (60 days) mouse. One sample was collected from spleen tissue (SP) and one from bone marrow (BM) and each sample was processed and sequenced two times independently resulting into four samples. Cell and transcript counting was performed and interpedently for each of the four samples using CellRanger (version 5.0.1) against a reference combining host and parasite sequences (Fig. 1B). Cells from the two replicates of each tissue were integrated using Seurat (v4.0) and the cells with detected *L. donovani* transcripts were annotated and found to cluster together in the UMAP projections (Fig. 1C). In the bone marrowBM and spleenSP samples, 1.47% and 3.16% cells were found to be parasitized respectively (Fig. 1D). Replicates from each tissue reported similar infection rates with bone marrowBM samples reporting 1.48% and 1.47% and spleenSP samples reporting 3.2% and 3.1% parasitized cells (Figure S1 and Table S1).

**Figure 1:**
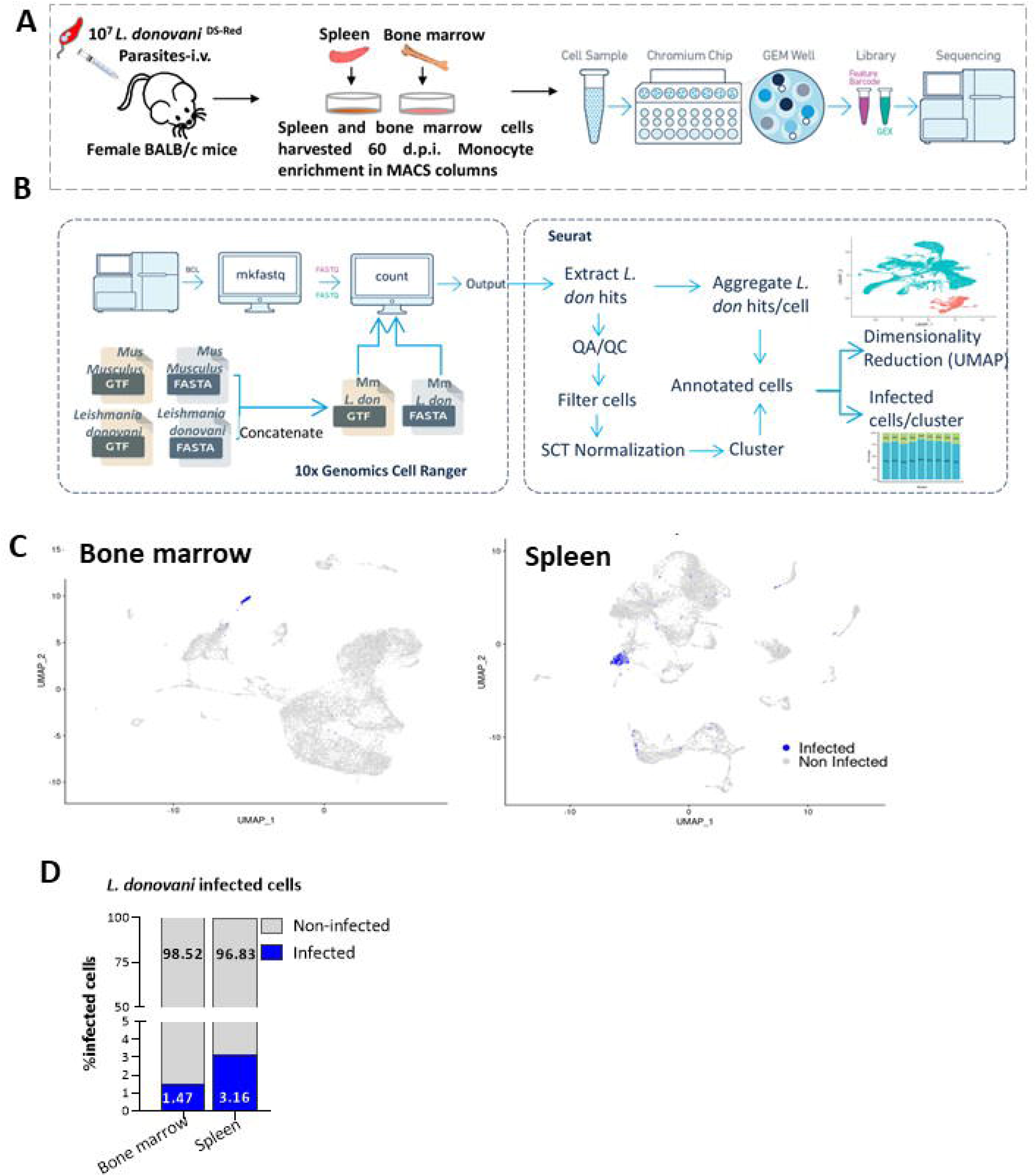
Schematic of the workflow of scRNA-Seq analysis. A) Spleen and bone marrow samples collected from mice chronically infected with *L. donovani* parasites were processed using 10X Genomics Chromium single cell RNA Seq (scRNA-Seq) technology. B) The schematic workflow for scRNA-Seq data processing. Base calling, read alignment and hit calling were performed using 10X Genomics CellRanger. *Mus Musculus* GRCm30 (Mm) and *L. donovani* (*L. don*) sequences were concatenated and the reads were aligned against a single reference. C) Uniform Manifold Approximation and Projection (UMAP) of ~14,000 cells from the spleen and ~14,000 cells from bone marrow where *L. don* infected cells are colored in blue. D) Bar graph showing the percentages of infected cells detected in each sample.

Cells were identified using cellranger’s adaptive algorithm resulting to a lower bound of 476 and 463 reads per barcode for the two spleen (SP) samples (Fig 2A-B) and 436 and 431 reads per barcode for the bone marrowBM samples (Fig 2C-D). The transcriptome of the host cells in all samples was not fully saturated (60%) reached typical levels for the technology, while the *L. donovani* transcriptome was not sufficiently saturated to perform any downstream analysis of the parasite’s transcriptome (Fig. 2A-D). A total of 18,197 and 18,111 genes were detected from the bone marrow replicates and 19,298 and 19,247 from the spleenSP replicates (Table 1, Table S1). Noticeably the mean number of reads per cell, the total number of cells and the number of UMIs mapped to the L .donovani reference were similar between the replicates (Table 1, Fig E, Table S1).

**Figure 2:**
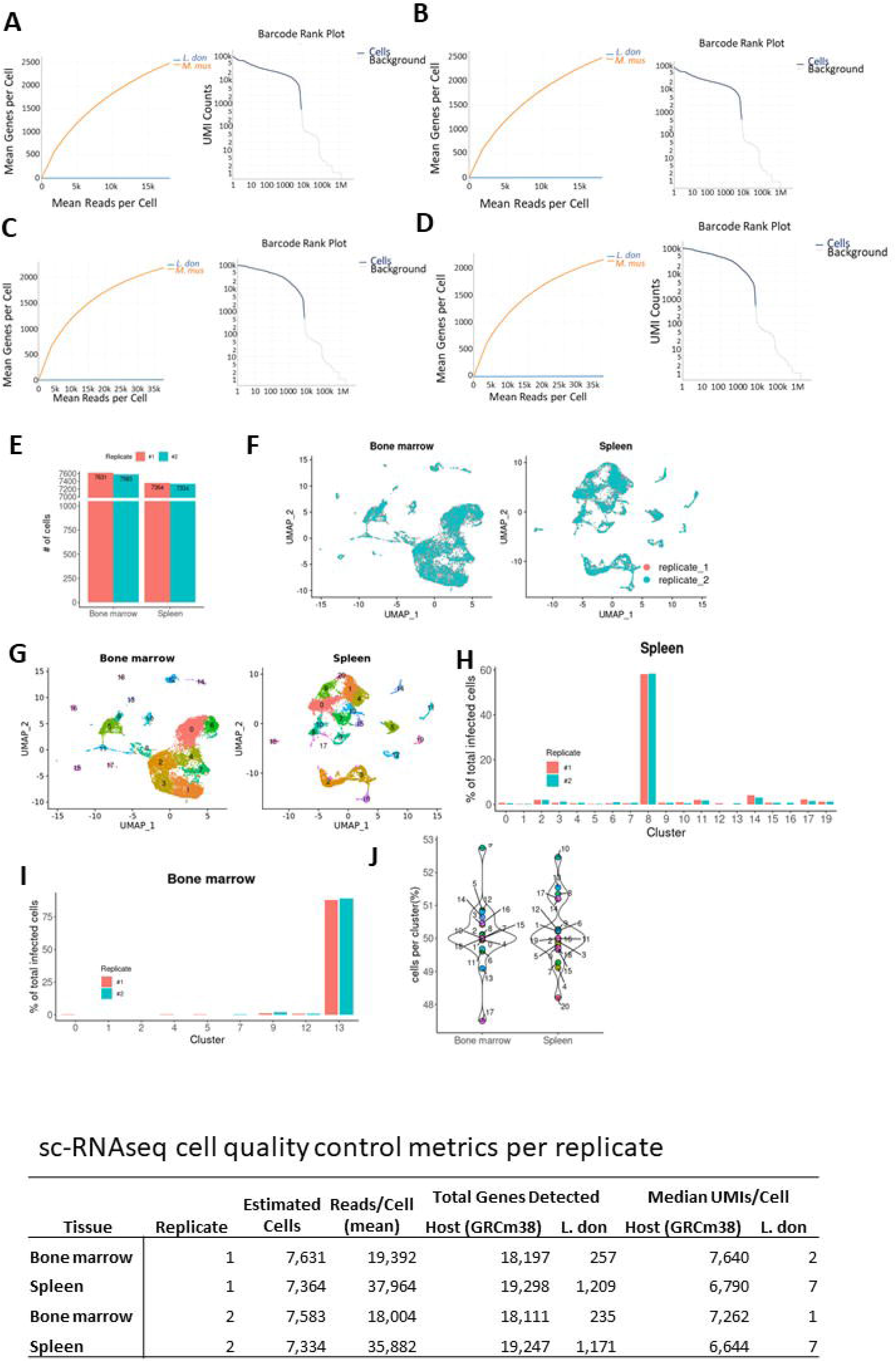
Bone marrow replicate 1 at 50% of the sequencing depth, yields 1805 genes and at 100% yields 2575, resulting in Δ50=30% saturation (A). Bone marrow replicate 2 at 50% of the sequencing depth, yields 1729 genes and 100% yields 2490, resulting in Δ50=31.6% saturatìon (B). Spleen replicate 1 at 50% of the sequencing depth, yields 1671 genes and 100% yields 2184, resulting in Δ50=23.5% saturation (C). Spleen replicate 2 at 50% of the sequencing depth, yields 1712 genes and 100% yields 2151, resulting in Δ50=20.5% saturation (D). In all four samples analyzed by cellranger, barcode ranking showed comparable cell counts. Histogram of total number of cells per replicate (E) and UMAP projection of integrated replicates from each tissue (F). G) UMAP projections of unsupervised clustered cells for BM and Spleen samples. Percentage of infected clustered cells for each replicate in Spleen (H) and Bone marrow (I) samples. J) Percentage of cells from replicate #1 of each clusterfor bone marrow and spleen samples.

**Table 1.**
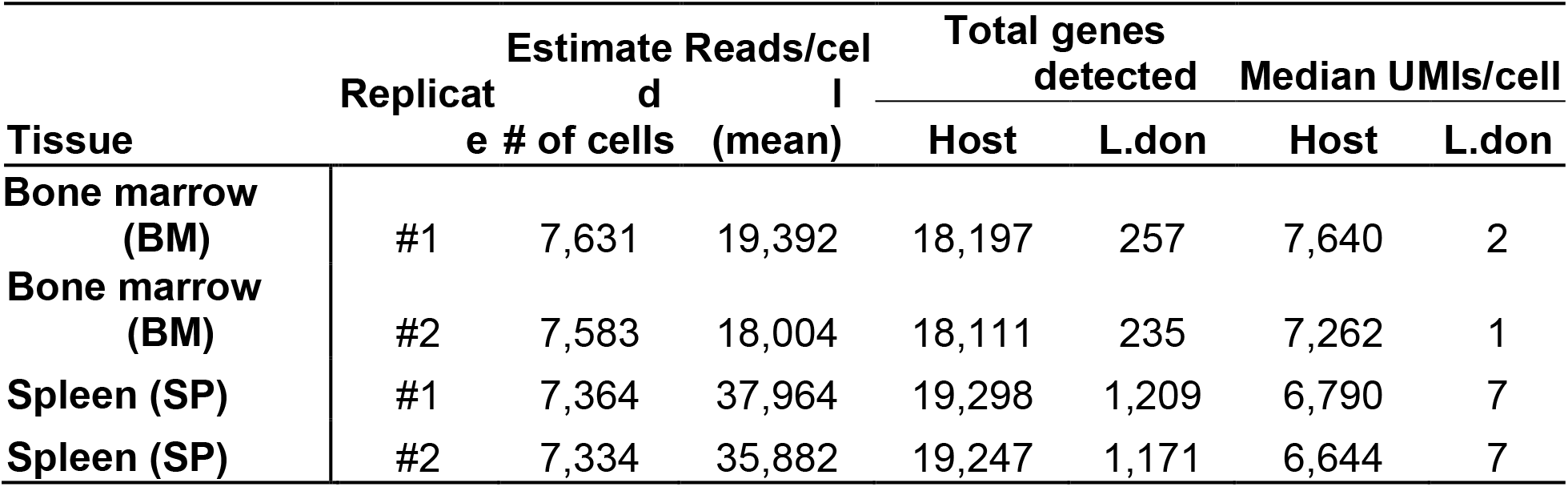
scRNAseq cell quality control metrics per replicate

### Populations of parasitized cells in BM and SP

Integration of the replicates from each tissue revealed no biases in the UMAP 2D projection with cell from one replicate overlapping with the cells of the other (Fig. 2F). Unsupervised clustering of the integrated cells revealed 19 and 21 clusters of a total 15,214 cells and 14,698 cells for the bone marrowBM and spleenSP samples respectively were integrated from the two spleenSP samples (Fig. G). Examination of total parasitized cells revealed the same distribution of clusters where infected cells were detected predominantly in one cluster (Fig. 1C), cluster #8 (~ 60%) and cluster #13 (~ 80%) in the spleenSP and bone marrowBM samples respectively (Fig. 2H and I). each cluster consists of cells from both replicates with no replicate contributing more than 53% cells in a cluster of either bone marrowBM or spleenSP samples (Fig. 2J).

Clusters from the integrated replicates were annotated into known cell populations from the CellMarker curated database (Zhang et al., 2019) using the list of marker genes of each cluster (Table S2, S3). In spleen, a total of 15 cell populations were identified, and 10 populations contained cells from a single cluster while the rest of the 11 clusters combined into 5 known populations (Fig. 3A). Similarly, in bone marrow, 11 cell populations were reported, out of which 6 where single cluster populations, while the rest of the 13 out of the 19 clusters were combined into 5 populations (Fig. 3D). The annotated cells were examined for replicate bias and no population was found to contain more than 53% of cells derived from one replicate: both for spleenSP (Fig. 3B) and bone marrowBM samples (Fig. E). Similarly, the distribution of infected populations from the total infected cells appeared to be unchanged between each replicate (Fig 3C and F). Inspection of the top 3 significant differentially expressed genes in each annotated cell population versus all other cells, revealed known markers such as Klrd1 (Cd94) for NK cells and Cd8b1 for CD8^+^ T cells in spleenSP (Fig. 3G and Table S4), and Flt3 (Mooney et al., 2017, Liu et al., 2013) for HSCs and Pf4 for megakaryocytes in bone marrowBM samples (Fig. 3H and Table S5).

**Figure 3:**
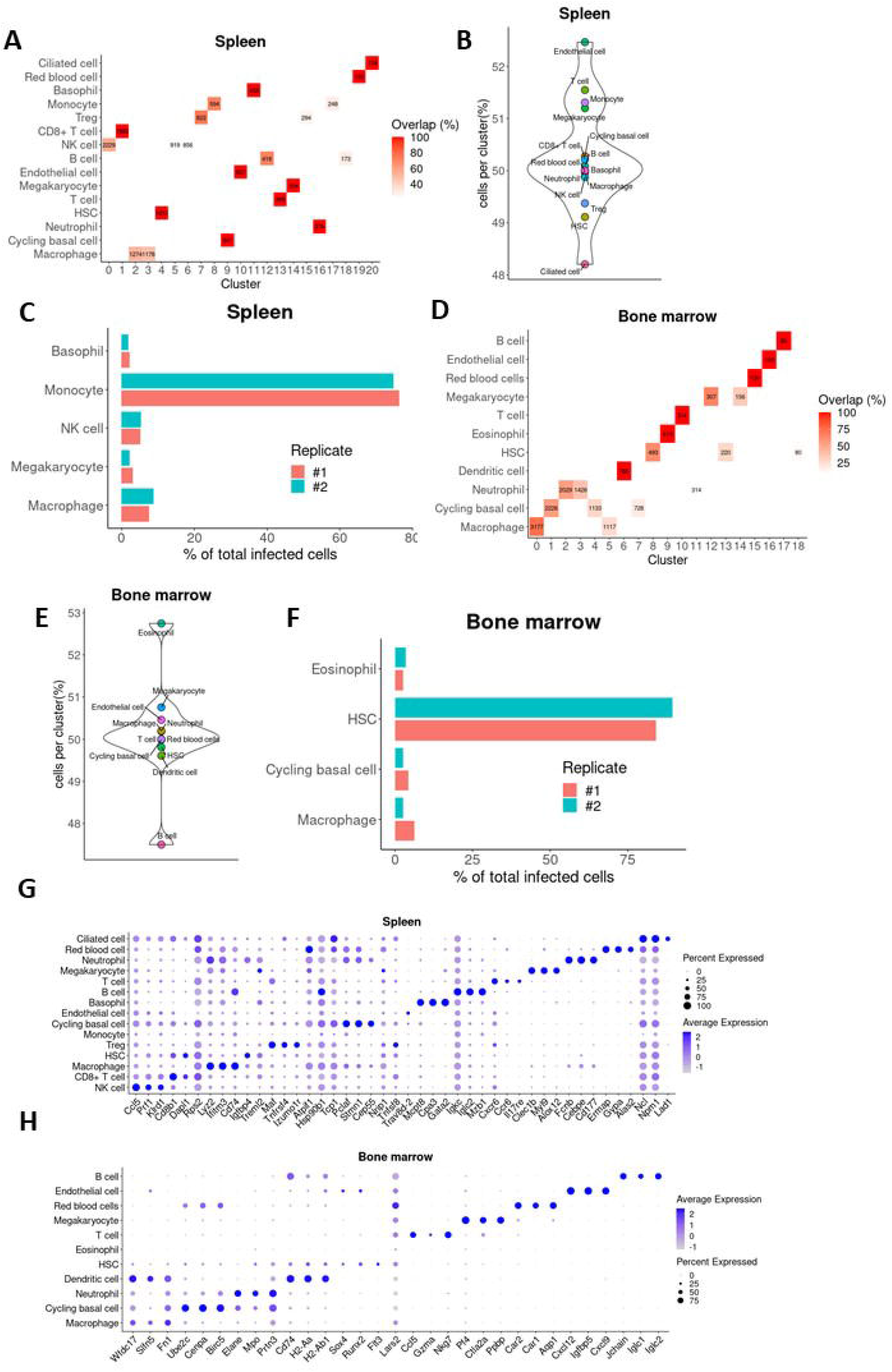
Annotation of cell populations. A) Confusion matrix between annotated cell populations and unsupervised clustering results from Spleen. B) Violin plot of the abundance of replicate #1 cells in the annotated cell populations from Spleen. C) Cell population distribution over the total infected cells per replicate from Spleen (C). Equivalent results from Bone marrow are shown (D-F). Expression levels of the top 3 genes from the list of markers used to annotate cell populations from Spleen (G) and Bone Marrow samples (H)

Annotation of cell populations in the UMAP projections showed that uninfected HSCs clustered farther from the infected HSCs, which was evidently the main population of parasitized cells in the bone marrowBM (Fig. 4A and C). In contrast, in the spleenSP samples, infected monocytes, which was the main parasitized population, were clustered together with uninfected monocytes. Although majority of the infected cells in the bone marrowBM samples were found in HSCs (95), 10 Macrophages, 8 Cycling basal cells and 7 Eosinophils were found to be parasitized (Fig. E). Similarly, in spleenSP samples, 9 Basophils, 38 Macrophages, 12 Megakaryocytes and 24 NK cells were detected to be parasitized along with the most prevalent parasitized populations, monocytes with 351 reported infected cells (Fig. 4F).

**Figure 4:**
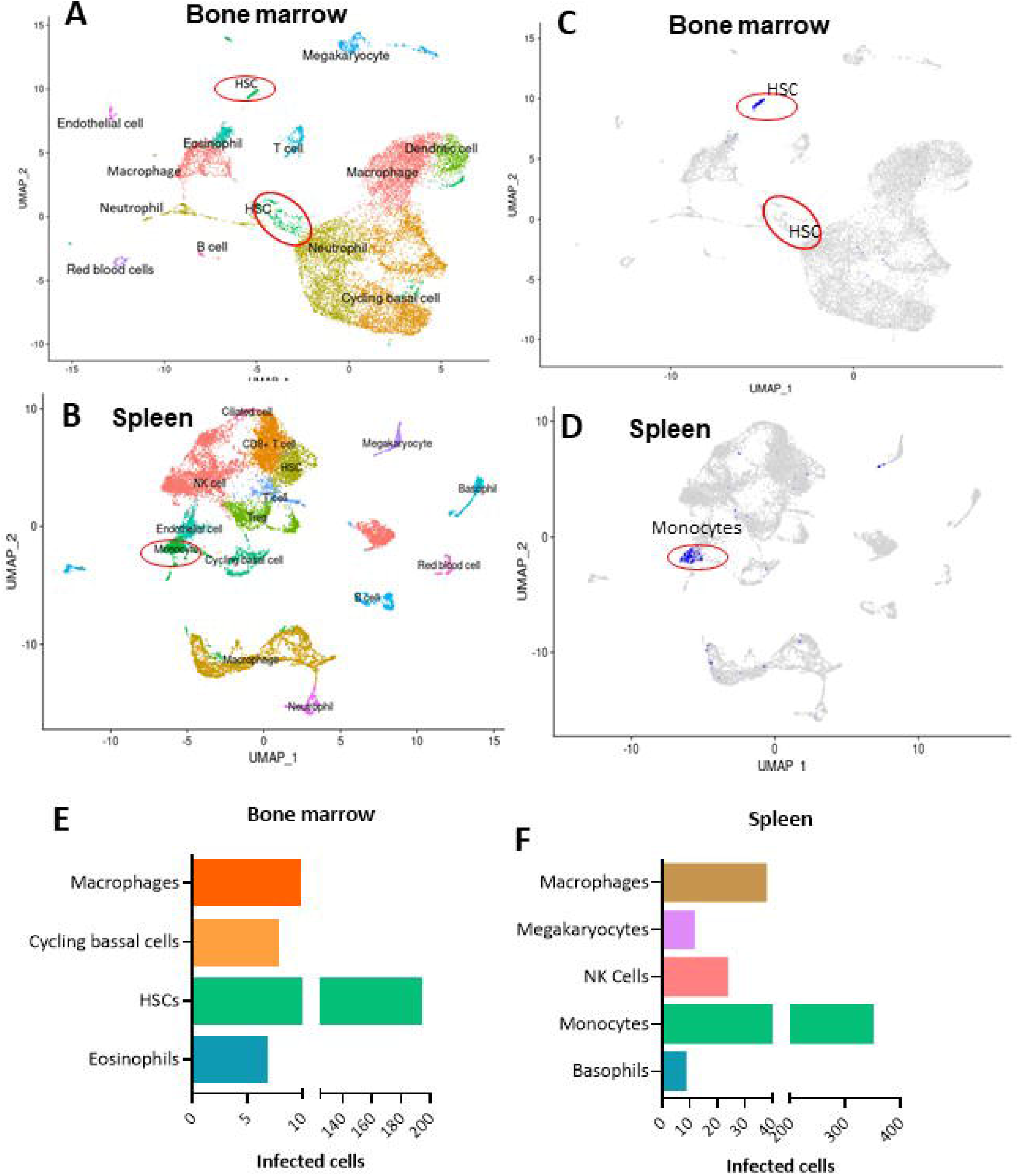
Single-cell transcriptomic UMAP embedding of A) bone marrow and B) spleen derived cells depicting color coded cell populations that were annotated based on CellMarker database. In bone marrow sample, C) two distinct hematopoietic stem cell populations are clustered differently where one constitutes the majority of *L. donovani* infected cells (a cluster of blue dots, shown in red circles). In spleen sample, D) a monocyte cluster was identified as the dominant population infected with *L. donovani* parasites (red circle). Distribution of infected cells across different populations in bone marrow (**E)** and **F)** spleen.

### scRNA-seq enables identification of differentially expressed transcripts in parasitized HSCs in BM and monocytes in spleen

Since *Leishmania donovani* transcripts were found in different frequencies in the host cells in different organs, we focused on HSCs and monocytes in bone marrow (BM) and spleen (SP) samples respectively. Splitting each population into two groups of parasitized and non-parasitized cells and performing a differential gene expression analysis revealed 602 and 1514 significantly differentially expressed genes in spleenSP and bone marrowBM respectively (Table S6 and S7). Interestingly, multiple genes, including Xist, Ptprc and Son, are present in the list of top 50 most significant genes (ranked by Bonferonni corrected p-value) from both spleenSP and bone marrowBM samples (Fig. 5A and C). Gene set enrichment analysis on KEGG pathways showed that pathways were mainly overrepresented in parasitized cells for both bone marrowBM and spleenSP samples (Fig. 5B and D and Table S8 and S9). Notably, Fcγ and CD93 receptors known to be involved in the phagocytosis of the parasite was found to be overrepresented in monocytes and HSCs.

**Figure 5:**
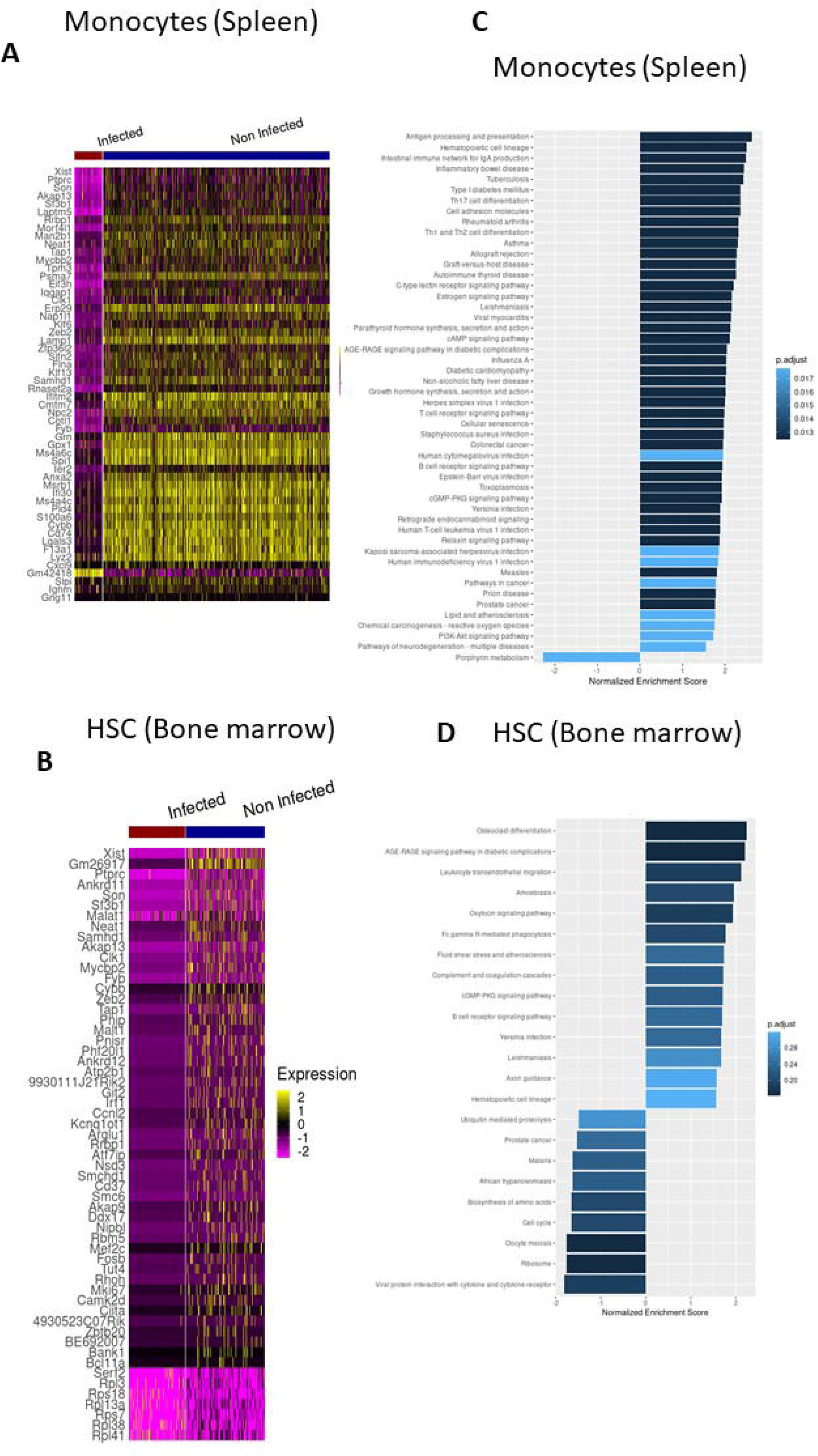
Differential gene expression in parasitized HSCs from Bone Marrow and Monocytes from Spleen. Heatmap of differentially expressed genes between the infected and non infected monocytes from the spleen (A) and HSCs from the bone marrow sample (B). Top 50 most significant, based on Bonferroni corrected p value, genes are shown. GSEA analysis showing select signaling networks enriched in *L. donovani* infected Monocytes (C) and HSCs (D).

### Infection of HSCs purified from mouse bone marrow

To confirm the results from scRNA-seq analysis that revealed HSCs in BM are the most frequent host cells for *L. donovani* parasites, we isolated hematopoietic progenitor cells from mouse BM and infected *in vitro* with virulent transgenic *L. donovani^RFP^* parasites. Flow cytometric analysis revealed that 40-50% of Lin- [Ter-119-CD11b-CD45R-CD5-CD3-Gr1-] c-Kit+Sca-1+HSCs indeed are infected as indicated by the RFP+ population (Fig. 6A). Analysis of *L. donovani*^RFP+^ HSCs revealed that almost all of the infected HSCs expressed Fcγ receptors CD16/32 (Fig 6B) and to some lesser extent CD93 (Fig 6C). A time course of infection revealed that *L. donovani*^RFP+^HSCs are detected as early as 24h after infection that progressively increased up to 72 hours postinfection (Fig. 6D).

**Figure 6:**
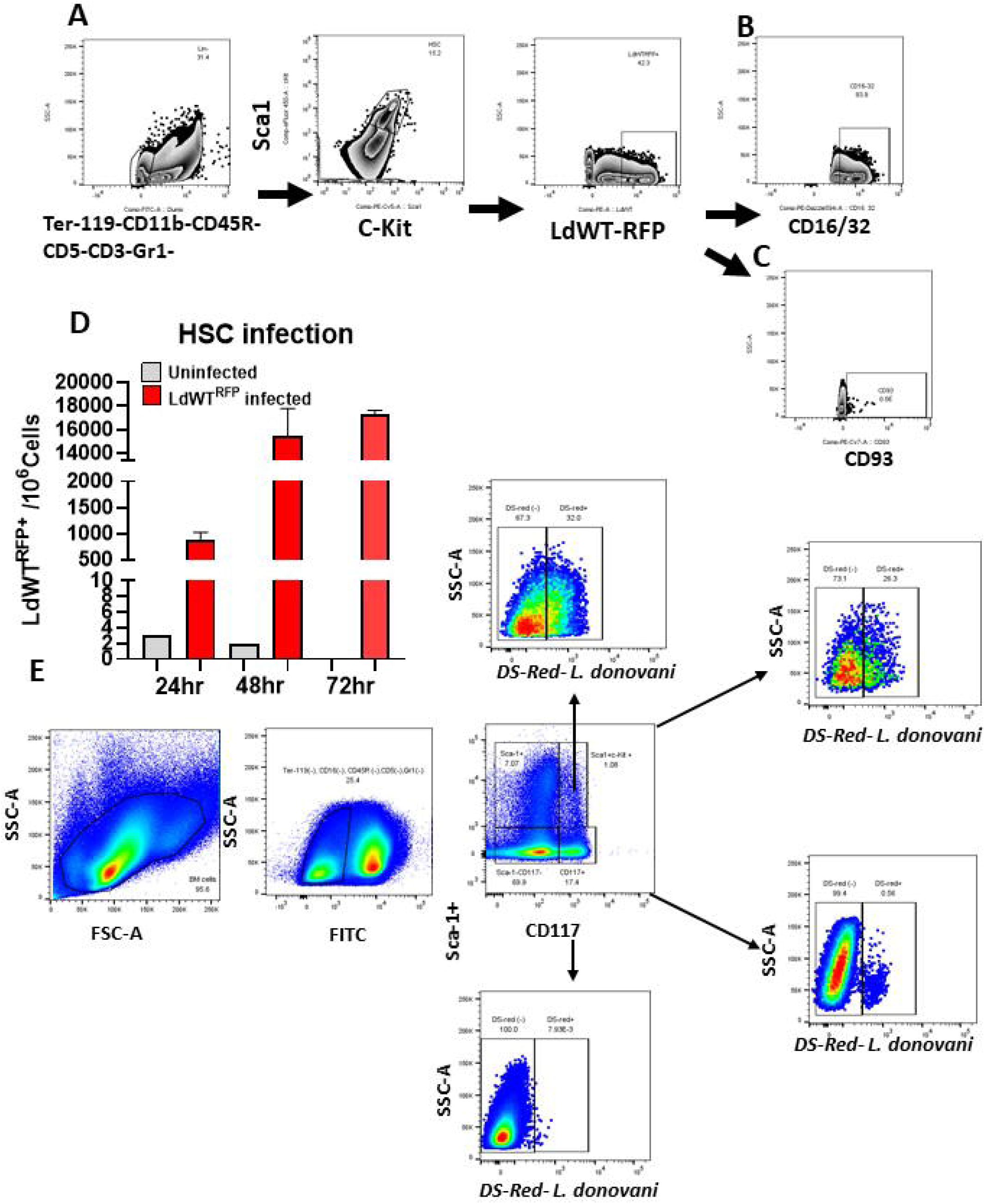
HSC infection of *L. donovani* in vitro. A) Hematopoietic progenitor cells were isolated from mouse bone marrow and were infected *in vitro* with transgenic LdWT^RFP^ parasites. Infected HSCs were identified by flow cytometry as Lin- [Ter-119-CD11b-CD45R-CD5-CD3-Gr1-] RFP+ cells. B) Consistent with our scRNA-Seq analysis, majority of the RFP+HSCs were also positive for CD16/32 indicating that uptake of the parasites might involve this receptor. C) Expression of CD93 was also observed on parasitized HSCs. D) Progressively increasing HSC infection with *L. donovani* parasites was observed in flow cytometric analysis. E) Identification of *L. donovani^RFP^* infected HSCs from a Balb/C mouse 9 months post infection.

### Identification of *L. donovani* infected HSCs in bone marrow from chronically infected mice

To verify the *in vitro* HSC infection results, we isolated BM from Balb/C mice 9 months post-infection with *L. donovani^DsRed^*. Consistent with scRNA-seq and in vitro experiments, flowcytometric analysis of BM cells revealed that Lin-c-Kit+Sca-1+ HSCs were infected with the parasites. Additionally, other populations of BM derived cells such as c-Kit-Sca-1+ cells were found to be infected as revealed by the DsRed expression.

## Discussion

Important problems remain to be solved towards the goals of control and elimination of human VL that causes significant mortality. The main hurdles that thwart the progress towards this goal are lack of understanding of the molecular basis of pathogenesis, persistent infections, drug resistance, incomplete cure that may cause PKDL Availability of a high throughput method to detect and annotate parasitized and bystander cells will open new ways of characterizing host-parasite interactions. Such a method could help identify rare cells that elude detection by conventional methods that employ bulk nucleic acid detection. Cataloguing of the infected cells in acute and chronic stages of *L. donovani* infection may enable targeting the sources of persistent infection and help identify mechanisms of drug resistance and immune protection in immunized or resistant individuals. Reliance on cell surface markers used in flow cytometry is inherently a limitation since it requires *a priori* knowledge of the cell types potentially infected and suitable reagents. Dual scRNA-seq method described here represents a breakthrough since it allows to agnostically examine the diversity of cell populations that may be infected with *Leishmania* parasites and to reconstruct the immune signatures that underlie the pathogenesis.

Analysis of SP samples by dual scRNA-Seq revealed that monocytes are the most frequently infected cell population. Previous studies in our laboratory have shown that inflammatory monocytes to be infected with *L. donovani* parasites in the SPs of chronically infected mice (Terrazas et al., 2017). Inflammatory monocytes develop in the BM and migrate to the site of infection in presence of IFN-γ produced by CD4^+^ T cells and develop into macrophages and dendritic cells (Romano et al., 2021). Identification of monocytes as the dominant population of infected cells in the SP by dual scRNA-Seq consistent with the previous literature in mouse model of chronic VL attests to the power of our method. Macrophages are the prototypical phagocytic cells infected with *Leishmania* parasites and thus widely studied. Imaging flow cytometry studies similarly showed the interplay between *L. donovani* parasites and macrophages in chronically infected mouse models (Terrazas et al., 2015). scRNA-Seq analysis based phenotyping of the parasitized and bystander monocytes and macrophages in the SP would reveal the role of these cells in pathogenesis. Unexpectedly our scRNA-Seq analysis revealed that NK cells, megakaryocytes and basophils to be parasitized in the SP in the chronic stage of infection. The phagocytic activity of NK cells has been reported in viral and bacterial pathogens (Abel et al., 2018), however to our knowledge uptake of *Leishmania* parasites has not been reported. Basophils are circulating cells that represent <1% of circulating leukocytes with a notable expression of IL-4 and IL-13 (Stone et al., 2010). Data on the role of megakaryocytes in *Leishmania* pathogenesis is sparse. The role of NK cells, basophils, and megakaryocytes in the splenic environment of chronic VL remains to be explored.

Analysis of BM samples by dual scRNA-Seq revealed unexpected results. Our results showed that hematopoietic stem cells (HSC) were the most numerous cell populations that were infected with *L. donovani* parasites. Previous studies indicated that HSCs and HSC like cells are unlikely to be infected with *L. donovani* parasites (Cotterell et al., 2000^a^, Cotterell et al., 2000^b^, Abidin et al., 2017, Hammami et al., 2017). BM has been studied since hematopoiesis is documented to be severely impacted in VL and other *Leishmania* infections (Mirkovich et al., 1986, Bandeira Ferreira et al., 2022). In vitro infection of HSCs purified from murine BM with transgenic *L. donovani* parasites that express RFP showed that HSCs were indeed infected with *L. donovani* parasites. HSCs were identified in our flow cytometric analysis as Ter-119-CD11b-CD45R-CD5-CD3-Gr1-Sca-1+c-Kit+ cells as was reported earlier (Kauffman et al., 2018, Khan et al., 2020, Pietras et al., 2015). Phagocytic potential of HSCs has been demonstrated in bacterial pathogens (Kolb-Maurer et al., 2002) and that HSCs quiescent under normal conditions are activated by inflammatory signals such as IFN-γ in chronic infections and undergo proliferation (Baldridge et al., 2010, Baldridge et al., 2011). Interestingly, parasitized HSCs showed an upregulation of FCγ receptor in our transcriptional analysis indicating that *L. donovani* parasites maybe using this receptor to gain entry into the HSCs. Consistent with this hypothesis, our flow cytometric analysis showed that almost all the *L. donovani*^DsRed+^ HSCs were also positive for the FCγ receptors CD16/32. Blocking studies in future will reveal the mechanism of entry of *L. donovani* parasites into HSCs. Recent studies showing the uptake of *L. donovani*/*L. infantum* parasites in human (Carvalho-Gontijo et al., 2018) and murine HSCs (Dirkx et al., 2022) further validates our results obtained by dual scRNA-Seq analysis. Our in vivo studies from Balb/C mice that has been infected with *L. donovani*^DsRed^ parasites for 9 months showed that Ter-119-CD11b-CD45R-CD5-CD3-Gr1-Sca-1+c-Kit+ HSCs remain infected with the parasites strengthens the hypothesis that HSCs maybe a source of persistent infection due to their longevity. Tracking of infected HSCs in bone marrowBM chimeras will test this hypothesis and reveal if *L. donovani* parasites remain dormant in HSCs with limited proliferation. Detection of eosinophils and cyclic basal cells as potentially parasitized population in BM was a surprising result in our analysis. Eosinophils have been studied in *L.amazonensis, L major* and *L. infantum* infections where they are recruited to the site of infection and show anti-microbial activities that helps in parasite clearance. Eosinophils have also been shown to possess antigen presenting activity and prime adaptive immunity (Rodriguez and Wilson 2014). Eosinophils have limited lifespan in tissues that ranges from 2-5 days however cytokines increase their survival in vitro to 14 or more days (Park et al., 2010). Presence of *L. donovani* parasites in short lived eosinophils and cycling basal cells presents intriguing possibilities. Does the presence of L donovani parasites in the shortlived cells such as eosinophils represent a transitory state of infection? What might be the contribution of these infected cells in the chronic stages of VL? Future studies addressing these questions will illuminate the pathophysiology of VL.

Recent discovery of trained immunity and its determinants in *M. tuberculosis* and BCG studies (Divangahi et al., 2021, Esaulova et al., 2021, Khader et al., 2019) showed that even though HSCs are not infected with these agents, the infection alters the epigenetic pathways in HSCs and such progeny acquire protection or pathogenic characteristics (Kaufmann et al., 2018, Khan et al., 2020). Application of dual scRNA-Seq will provide a powerful method to address epigenetic reprogramming in *L. donovani* infection bone marrow chimera studies to identify mechanisms of protection. Application of additional analytical methods such as ATAC seq in parasitized cells (Perkel 2021) will help track cell fate decisions and advance our understanding of hematopoietic defects in human VL.

Previously, bulk RNAseq has been employed to explore transcriptional changes in the cells of infected hosts as opposed to non-infected animals as well as the transcriptome of the parasite (Forrester et al., 2022). But due to the intrinsic limitations of the bulk nature of the technology, the relation between the differentially expressed genes and the parasitized cells is a task yet to be solved. Analytical methods that deconvolve bulk RNA-seq data to the level of cell populations are available in the literature but their accuracy is significantly below that of singe cell technologies, and they depend on prior knowledge of the populations present in the sample, as well as the transcriptional profile of such populations (Gong and Szustakowski 2013). Many public resources, such as ImmGen and the Human Cell Atlas, provide the required data for the desired cell populations, yet there is no information available for parasitized cell types, which would be required to deconvolve infected from uninfected cells. Logically, past attempts of deconvolution have compared results between infected and uninfected tissue (Dirkx et al., 2022). However, deconvolution of cell populations derived an infected host using prior knowledge of cell types from uninfected host remains a challenge of this approach. On the other hand, acquiring single level information on the parasitized status of the cells among the same population allowed us to investigate potential receptors associated with the phagocytosis of the parasite. These observations were borne out by the in vitro infection of purified HSCs with transgenic *Leishmania donovani* parasites followed by flow cytometric analysis. Future studies with knockout mouse models or blocking antibodies will provide definitive evidence of the involvement of these receptors in HSCs in acquiring Leishmania parasites.

Despite the major increase in resolution offered by single cell RNA-seq, the technology, hence our approach, has limitations. The ability to study heterogenous populations and investigate the transcriptome of each cell individually is compensated by a considerably decreased saturation of their transcriptional profile compared to what is routinely observed in bulk RNA-seq, leading to dropouts. Dropout is the phenomena when lowly or moderately expressed genes are not detected in some cells. Given the low number of detected *L. donovani* transcripts it is reasonable to expect that the number of true infected cells in our analysis maybe higher. Another limitation we observed, similar to the dropout effect, is the low depth of coverage on *L. donovani’s* transcriptome. Due to this shortcoming, we were unable to follow downstream analysis and gain further insight of the parasite’s genes and pathways preferentially involved in each infected cell population.

Availability of high quality reference genomes of *L. donovani* (GCF_000227135.1) enabled clear cut identification of parasitized cells when *L. donovani*. Detection of host transcripts was relatively straightforward in our method. However, detection of *L. donovani* remains a challenge. Performing additional rounds of sequencing only resulted in diminishing benefit in enhancing parasite transcript recovery as shown in our saturation curves. It is conceivable that including additional barcodes targeting the splice leader sequences present in all mature transcripts of *Leishmania*, or technologies with higher per cell throughput, may allow improved capture of parasite transcripts. Our method is tunable to incorporate additional inputs from parasite transcriptome.

The method described in this paper to reliably detect and annotate *Leishmania* parasite infected cells in mouse models can be extended to the analysis of human bone marrow aspirates with relative ease. The interactome of parasite and host cells will be amenable to enquiry with unprecedented resolution with our method. Probing complex problems such as persistent parasitic infection, immune mechanisms that enable low level chronic infection, failure of anti-leishmanial drugs to eliminate the reservoirs of infection, effect of *Leishmania* infection on hematopoiesis, anemia and trained immunity that remain elusive due to lack of precision methods can now be investigated.

